# Profiling the *in vitro* and *in vivo* activity of streptothricin-F against carbapenem-resistant Enterobacterales: a historic scaffold with a novel mechanism of action

**DOI:** 10.1101/2021.06.14.448463

**Authors:** Kenneth P. Smith, Yoon-Suk Kang, Alex B. Green, Matthew G. Dowgiallo, Brandon C. Miller, Lucius Chiaraviglio, Katherine A. Truelson, Katelyn E. Zulauf, Shade Rodriguez, Roman Manetsch, James E. Kirby

**Affiliations:** Department of Pathology, Beth Israel Deaconess Medical Center, Boston MA; Harvard Medical School, Boston MA, USA; Department of Chemistry and Chemical Biology, Northeastern University, 360 Huntington Ave, Boston, MA 02115, USA; Department of Pharmaceutical Sciences, Northeastern University, 360 Huntington Ave, Boston, MA 02115, USA; Center for Drug Discovery, Northeastern University, Boston, MA 02115, USA

## Abstract

Streptothricins are components of the natural product, nourseothricin; each containing identical streptolidine and gulosamine aminosugar moieties attached to varying numbers of linked β-lysines. Nourseothricin was discovered by Waksman and colleagues in the early 1940’s, generating intense interest because of excellent Gram-negative activity. However, the natural product mixture was associated with toxicity, and subsequent exploration was limited. Here, we establish the activity spectrum of nourseothricin and its main components, streptothricin-F (S-F, one lysine) and streptothricin D (S-D, three lysines), purified to homogeneity, against highly drug-resistant, carbapenem-resistant Enterobacterales (CRE). The MIC_50_ and MIC_90_ for S-F and S-D were 2 and 4 µM, and 0.25 and 0.5 µM, respectively. S-F and nourseothricin showed rapid, bactericidal activity. S-F and S-D both showed approximately 40-fold greater selectivity for prokaryotic than eukaryotic ribosomes in *in vitro* translation assays. There was >10-fold *in vitro* selectivity of S-F compared with S-D on LLC-PK1 and J774 cell lines. *In vivo*, delayed renal toxicity occurred at >10-fold higher doses of S-F compared with S-D. Substantial treatment effect of S-F in the murine thigh model was observed against the otherwise pandrug-resistant, NDM-1-expressing *Klebsiella pneumoniae* Nevada strain at dosing levels without observable or minimal toxicity. Resistance mutations obtained in single ribosomal operon *E. coli* identify novel interactions with 16S rRNA helix 34, i.e., C1504A and A1196G/C conferred high level resistance to nourseothricin. Based on promising, unique activity, we suggest that the streptothricin scaffold deserves further pre-clinical exploration as a potential therapeutic for the treatment of CRE and potentially other multidrug-resistant, gram-negative pathogens.

**IMPORTANCE:** Streptothricins are a historic class of antibiotics discovered by Waksman and colleagues in the 1940’s. Toxicities associated with the streptothricin natural product mixture, also known as nourseothricin, discouraged further development. However, we found that a component of nourseothricin, streptothricin-F, retained potent activity against contemporary carbapenem-resistant Enterobacterales with significant selectivity in *in vitro* and *in vivo* assays. This included demonstration of rapid bactericidal activity *in vitro* and substantial therapeutic effect in the murine thigh model against a pandrug-resistant *Klebsiella pneumoniae* isolate at non-toxic concentrations. Through resistance mutation analysis, we identified helix 34 of 16S rRNA in the prokaryotic ribosome, and specifically bases C1054 and A1196, as critical for streptothricin’s activity. The mechanism of action is distinct from other known translation inhibitors. Based on promising and unique activity, we believe the streptothricin scaffold deserves further pre-clinical exploration as a potential therapeutic for the treatment of CRE and potentially other multidrug-resistant, Gram-negative pathogens.

## INTRODUCTION

The rapid emergence of antimicrobial resistance presents a significant challenge for treatment of bacterial infections. Carbapenem-resistant Enterobacterales (CRE) are of particular concern. The gram-negative cell envelope presents a formidable barrier to potential antimicrobial compounds based on a double cell membrane permeability barrier and efflux pumps. Almost all approved gram-negative antimicrobials that can overcome this barrier are natural products or synthetic or semi-synthetic derivatives of natural products. Unfortunately, small molecules commonly available in high-throughput screening libraries rarely share similar physicochemical properties associated with gram-negative penetrance and activity (1). As such, high throughput screening efforts using synthetic chemical libraries with rare exceptions have been non-productive (1). As a result, there is a significant antimicrobial discovery void.

Moreover, there is little doubt that resistance will emerge to agents currently in the pipeline. It is already known for example, that plazomicin, an aminoglycoside engineered to avoid drug-modification-based resistance mechanisms, is ineffective against strains with ribosomal methylase-based resistance mechanisms found in a large proportion of NDM-1 CRE strains (2, 3). Concerningly, these NDM-1 strains, highly endemic in East Asia (in particular India) (4-8), are emerging in the US, exemplified by the report of a lethal NDM-1 CRE infection in Nevada that was resistant to all available agents (4). We are therefore clearly in need of several new gram-negative agents that are unique in terms of antimicrobial class and potential vulnerabilities, and which can diversify our antimicrobial therapeutic portfolio (9).

Here, we further consider the properties of a historic antibiotic scaffold. Streptothricin was originally isolated by Waksman and Woodruff in 1942 from a soil *Actinomyces* (10). It generated initial excitement based on activity against gram-negative organisms and *Mycobacterium tuberculosis* (11). It was also noted to completely cure *Brucella abortus* infection in guinea pigs (12) and otherwise lethal *Salmonella paratyphi* B infection in mice (13). Based on these attributes, it went into fermentative production at Merck. However, in a limited human trial (details not published) (14), the natural product streptothricin induced reversible kidney toxicity, and, therefore, was not further pursued as a therapeutic.

Studies forty years ago using *in vitro* translation systems determined that the natural product inhibited prokaryotic translation and induced substantial ribosomal miscoding activity, similar to aminoglycosides (15). Notably, however, it did not inhibit translation in rat liver extracts, suggesting prokaryotic selectivity.

Streptothricin is now known to be a natural product mixture, often referred to as nourseothricin. For clarity, we will henceforth refer to the natural product mixture as nourseothricin and individual components by their specific streptothricin designation. Each streptothricin in the nourseothricin natural product mixture consists of three linked components, a streptolidine, a carbamoylated gulosamine moiety, and a β-lysine homopolymer of varying length. The individual streptothricins are designated by letter suffixes (A-F and X) based on the number of lysine residues with streptothricin F (S-F) and streptothricin D (S-D) having one and three lysine moieties, respectively (see Fig. 1).

**Figure 1.**
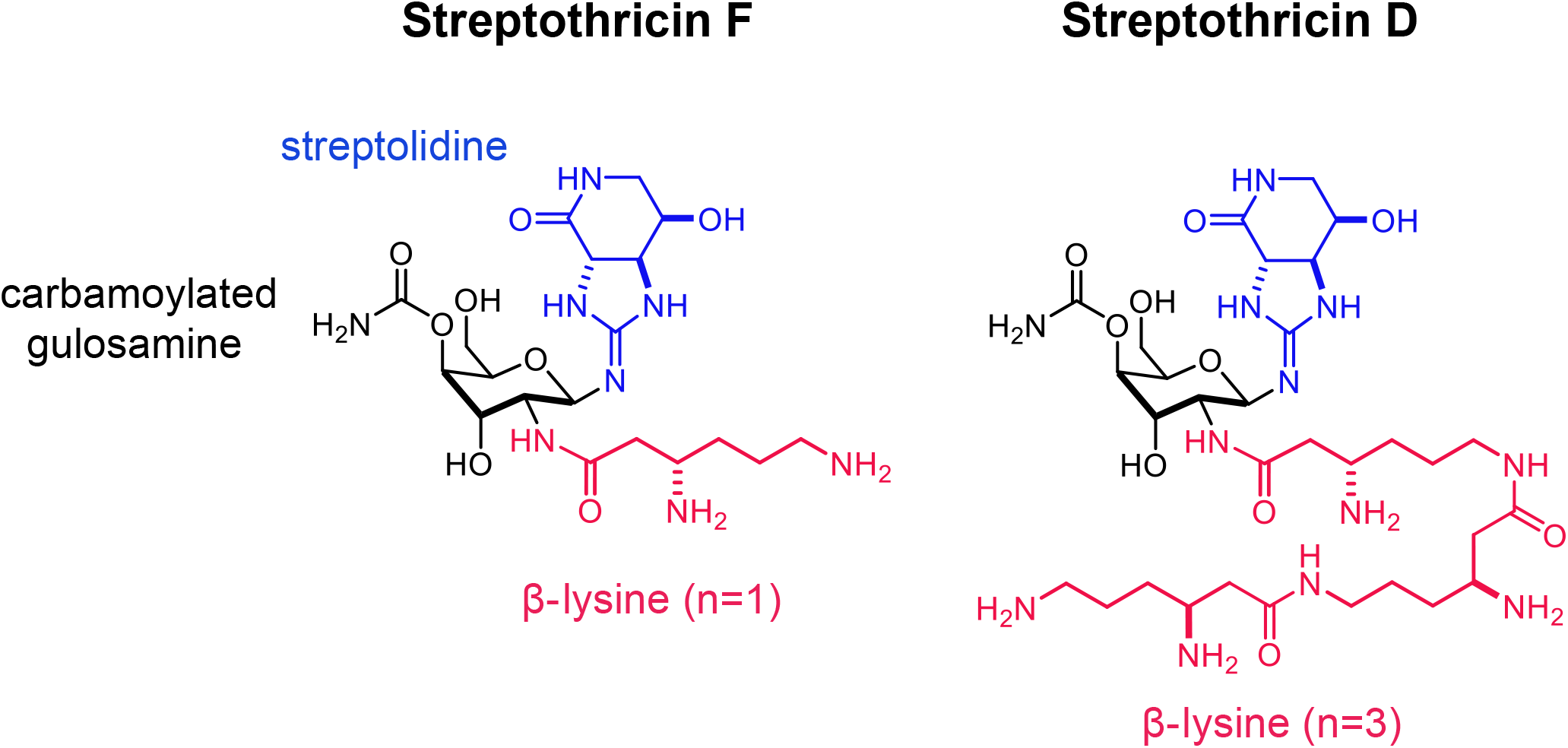
Structure of streptothricin F and streptothricin D. Streptothricins share streptolidine and a carbamoylated gulosamine sugar moieties. They are distinguished by differing numbers of β-lysines attached end-to-end through amide bonds to the 6-amino groups. Nourseothricin is the natural product mixture of several streptothricins, predominantly streptothricin F (one β-lysine) and streptothricin D (three β-lysines). Acetylation of the β-amino group blocks activity and is the major known mechanism of antimicrobial resistance to streptothricins.

Initial studies on toxicity in animals from the 1940’s used only semi-quantitative activity measures and impure compound with large doses presumably exceeding 100 mg/kg (16) and are therefore problematic to interpret. More recent studies in mice with fractionated streptothricins showed that toxicity varies with the length of the poly-β-lysine chain: specifically, a murine LD_50_ of 300 mg/kg for S-F with one lysine compared to an LD_50_ of ∼10 mg/kg for S-D and S-C with 3 and 4 β-lysines, respectively (16, 17). For comparison, the murine LD_50_ is 52 mg/kg i.v. for gentamicin (18); 40 mg/kg i.p. for colistin methanesulfonate (19); and 260 mg/kg i.p. for tobramycin (20). Streptothricin’s delayed toxicity towards renal proximal convoluted tubules, only demonstrated histologically in the literature for S-C in rats (21), has not been fully explained at a mechanistic level (16). Of note, S-C at 40 mg/kg, well above the lethal dose, did not cause liver damage as assessed by serum aspartate aminotransferase (AST) and alanine aminotransferase (ALT) enzymatic assays (21), highlighting kidney damage as the primary limiting toxicity.

Based on potentially insufficiently explored therapeutic promise for streptothricins, we sought to further characterize the properties of streptothricin-F and streptothricin-D, the major constituents of nourseothricin, purified to homogeneity using modern techniques and authenticated by several complementary state-of-the-art methods.

This analysis included activity spectrum studies against CRE, time-kill studies, prokaryotic and eukaryotic *in vitro* translation inhibition assays, maximum tolerate dose studies with histopathological analysis of renal pathology, and therapeutic efficacy testing in the murine thigh model. Taken together, these studies establish streptothricin-F as a natural product scaffold with what we believe are compelling properties for future medicinal chemistry exploration. To this end, we have separately developed a diversity-enabling total synthesis for S-F, which we will report elsewhere.

## RESULTS

### Activity spectrum studies

Our original studies with streptothricins were motivated by the observation that a nourseothricin resistance cassette served as a consistently tractable genetic marker in otherwise highly drug-resistant gram-negative pathogens (22, 23). This led to the question of whether nourseothricin and the component streptothricins had more general activity against multidrug-resistant gram-negative pathogens such as CRE. Streptothricin-F (S-F), as noted above, was observed to have much lower toxicity in mice and rats and therefore was of particular interest. We, therefore adapted a previously reported method to purify S-F (one β-lysine) and S-D (three β-lysines) to homogeneity using Sephadex chromatography (17). Confirmation of purity and precise molar content was determined by elemental analysis, NMR, and LC-MS as described in Supplementary Materials and Methods, Figs. S1-S3, and Table S1, S2. Purified S-D was available in much lower amounts than purified S-F. As the primary focus of our studies was S-F (infra vide), we therefore only tested S-D selectively in experiments.

Using broth microdilution minimal inhibitory concentration (MIC) testing, we determined the MIC of S-F, S-D, and nourseothricin for a large group of multidrug-resistant carbapenem-resistant Enterobacterales. These included strains expressing metallo- and/or serine carbapenemases and the pandrug-resistant, NDM-1 expressing *Klebsiella pneumoniae* Nevada strain, AR-0663 (4). For these CRE (n=39), MIC_50_ and MIC_90_ for S-F, S-D and nourseothricin, respectively were 2 and 4 (range 1-4) µM; 0.25 and 0.5 (range 0.25-2) µM; and 0.5 and 1 (range 0.25-2) µM. The average MIC was 5.6-fold greater for S-F than for S-D and 4.2-fold greater for S-F than for nourseothricin. Data for individual isolates are listed in Table S3.

An LC-MS analysis of commercially purchased nourseothricin showed a composition of 65.5% S-F; 29.6% S-D, and 4.9% S-E (2 β-lysines). Therefore, the somewhat enhanced activity of S-D (mean MIC 0.44 µM) compared with nourseothricin (mean MIC 0.58 µM) likely reflects a combinatorial effect of the more potent S-D and less potent S-F, as well as contributions from other minor streptothricin congeners in the nourseothricin natural product mixture.

### *In vitro* time-kill analysis

Bacterial killing by unfractionated, crude preparations of streptothricins was previously noted by Waksman and Woodruff for *E. coli* and *Micrococcus lysodeikticus*, after 17 and 65 hours of exposure without reference to an underlying MIC (10). To characterize potential bactericidal activity, we therefore performed time-kill analysis for S-F and nourseothricin against the pandrug-resistant CRE *Klebsiella pneumoniae* Nevada strain (AR-0636). Notably, we observed rapid bactericidal activity for both within two hours of exposure at 4X MIC without regrowth (Fig. 2). Therefore, bactericidal activity for a highly resistant CRE strain was observed using current methods.

**Figure 2.**
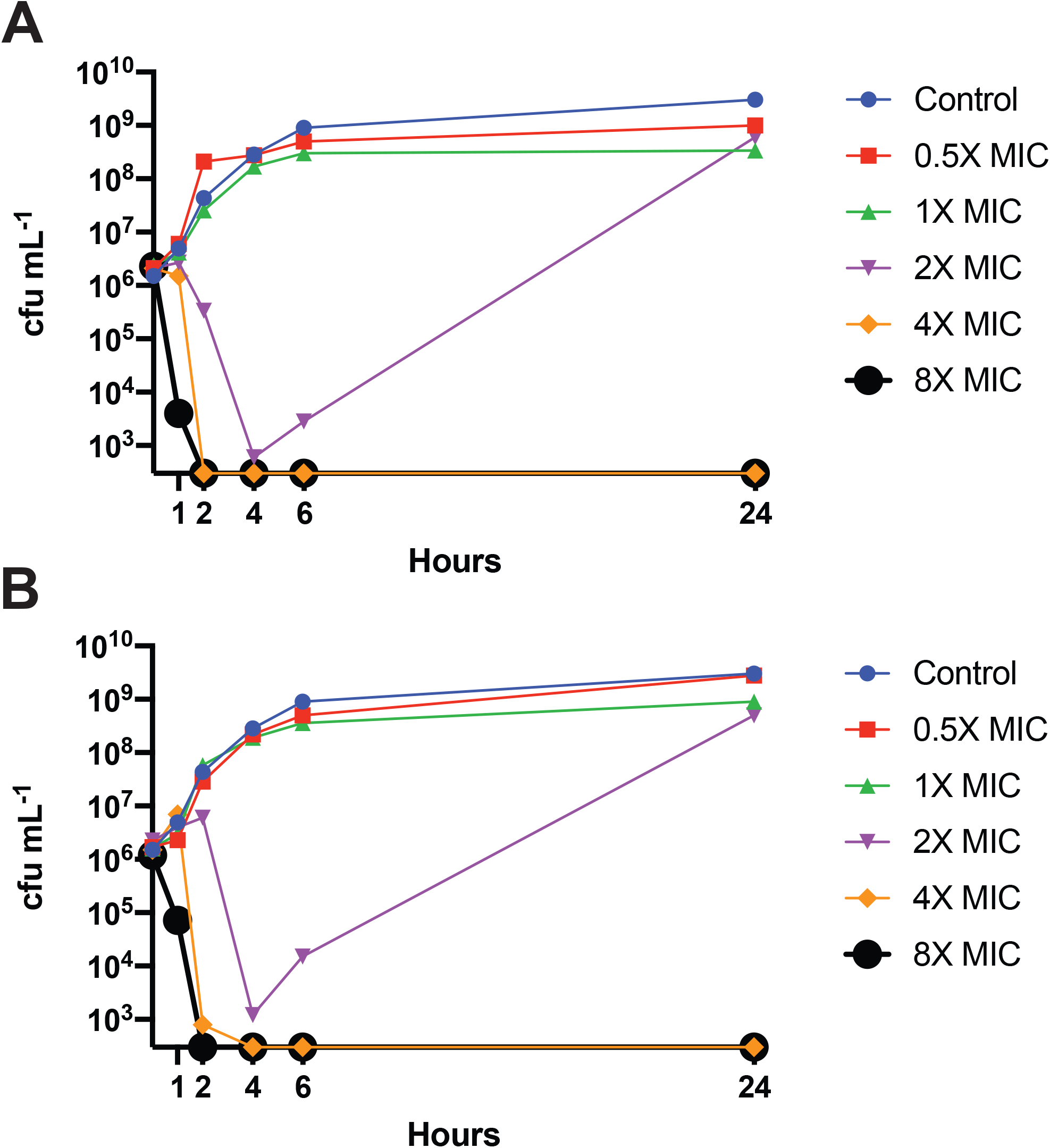
Rapid bactericidal activity against the *Klebsiella pneumonie* Nevada strain. **(A)** Nourseothricin MIC 0.25 µM. **(B)** Streptothricin-F MIC 1 µM.

### In vitro translation assays

We developed and validated assays to measure *in vitro* action on ribosome translation (i.e., independent of bacterial permeability and efflux effects) using commercially (NEB) available prokaryotic (*E. coli*) and eukaryotic (rabbit reticulocyte) *in vitro* coupled transcription-translation systems with readout using nanoluciferase gene constructs (see Fig. S4). In this analysis, we found that both S-F and S-D were approximately 40-fold more selective for prokaryotic than eukaryotic ribosomes based on IC_50_ values determined from Hill-Slopes of dose-response curves (Fig. 3). Interestingly, S-D was approximately 10-fold more potent than S-F on a molar basis, helping to explain the lower MIC’s noted for the former. However, the difference in translational inhibition was greater than the difference in mean MIC for S-F and S-D. We, therefore, considered whether S-F might have enhanced bacterial penetrance. However, the activity of S-D and S-F were not changed in isogenic *E. coli lptD* and *tolC* mutants (data not shown) suggesting that the outer membrane permeability and efflux pump activity as assessed by these measures were not significant barriers to S-F and S-D penetration.

**Figure 3.**
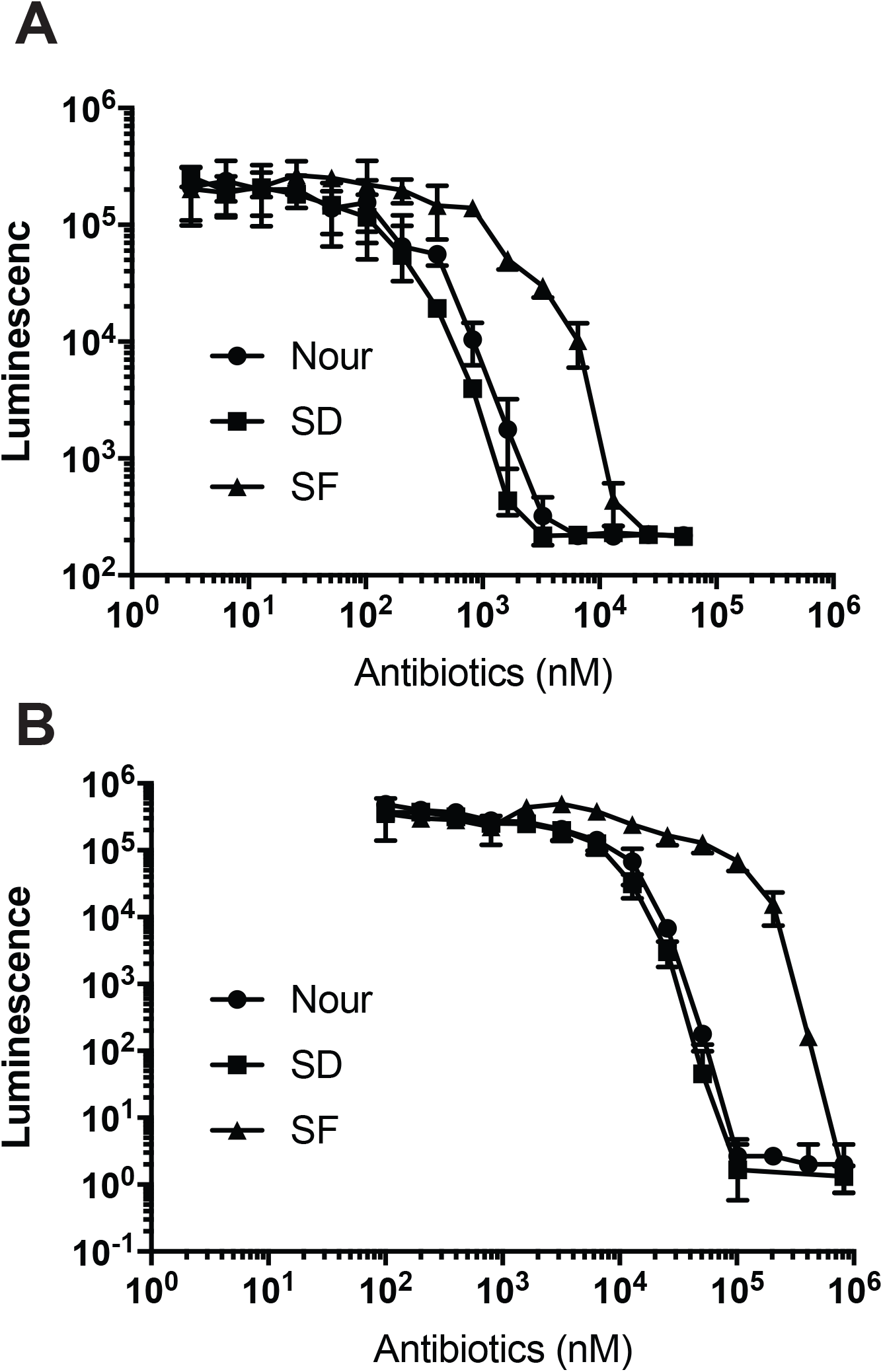
Inhibition of prokaryotic and eukaryotic translation. **(A)** Inhibition of prokaryotic in vitro translation using coupled *in vitro* transcription-translation extracts with readout from a nanoluciferase reporter. **(B)** Inhibition of eukaryotic in vitro translation using coupled *in vitro* transcription-translation extracts with readout from a nanoluciferase reporter. S-F, S-D and nourseothricin (nour) data represent mean and standard deviation from three independent experiments.

### Cytotoxicity

Eukaryotic cell cytotoxicity was determined against J774A.1 macrophage and LLC-PK1 proximal tubule kidney cells line during a five-day incubation. Cytotoxicity was measured using a previously established SYTOX Green exclusion, real-time fluorescence assay (24, 25). Notably CC_50_ for these cell lines was > 10X higher for S-F than S-D (Fig. 4). Cytotoxicity was also relatively delayed for S-F compared with S-D, first apparent on day 2 of incubation, with more prominent cytotoxicity for both S-F and S-D noted after five days of incubation. The delay in observed cytotoxicity was more pronounced for LLC-PK-1 in which cytotoxicity for S-D and S-F were not appreciable until day 3, and of borderline magnitude for S-F at day 5 (Fig. 4). Notably, the lowest observable adverse effect level for S-F during prolonged 5-day incubation was 32 µM, substantially above the MIC_99_ of 4 µM for CRE.

**Figure 4.**
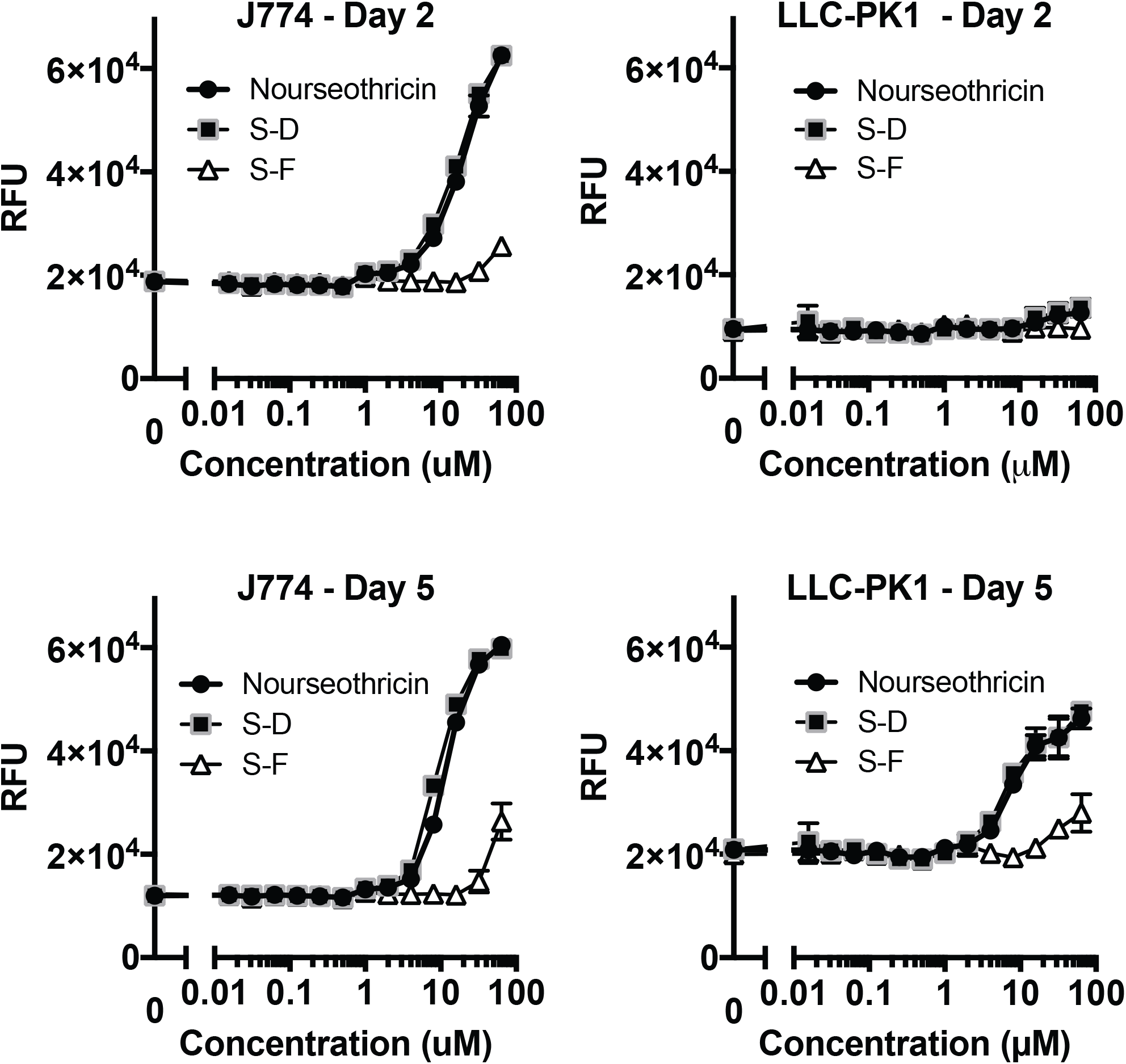
Cytotoxicity of streptothricins against mammalian cell lines. J774 macrophages and LLC-PK-1 renal epithelial cells were treated with two-fold doubling dilutions of nourseothricin, S-D and S-F for up to 5 days in the presence of SYTOX-Green. SYTOX-Green is a cell membrane-impermeant nucleic acid binding dye that fluoresces on binding to nuclear DNA. It therefore provides a real-time readout of eukaryotic cell membrane permeabilization associated with cell death that can be continuously monitored through fluorescence measurements. Cytotoxicity was minimal to absent after a single day incubation but increased on subsequent days. Nourseothricin and S-D effects were essentially indistinguishable. S-F toxicity was only observed at molar concentration at least 10-fold greater than S-D beginning at 32 μM, significantly above MIC ranges observed in activity spectrum analysis. Each data point represents mean and standard deviation for assays performed in quadruplicate.

### Maximum tolerated dose

To identify the single-dose maximum tolerated dose (MTD) of S-F and nourseothricin, CD-1 mice were injected intraperitoneally (IP) with ascending doses of 0, 50 100, 200, and 400 mg/kg S-F and 5, 10, 20 and 50 mg/kg nourseothricin, respectively (n=4 per dose). Nourseothricin was used as a surrogate for S-D, based on the difficulty of isolation of the latter in quantities needed for the experiments. Over the next 72 hours, no signs of distress were observed with dosing up to 200 mg/kg of S-F and 20 mg/kg of nourseothricin, respectively. Two mice at 400 mg/kg of S-F and 1 mouse at 50 mg/kg nourseothricin died, or became moribund and were euthanized. Histology of kidneys was specifically examined for all mice based on described delayed renal toxicity in rats for the streptothricin natural product. For S-F, proximal convoluted tubular damage was noted sporadically at 100 mg/kg and more diffusely at ≥ 200 mg/kg. For nourseothricin proximal tubular damage was pronounced at ≥10 mg/kg. Glomeruli, distal tubules and vasculature were spared (Fig. 5). Therefore, the maximum dose without observed pathological effect was 50 mg/kg for S-F and 5 mg/kg for nourseothricin, respectively.

**Figure 5.**
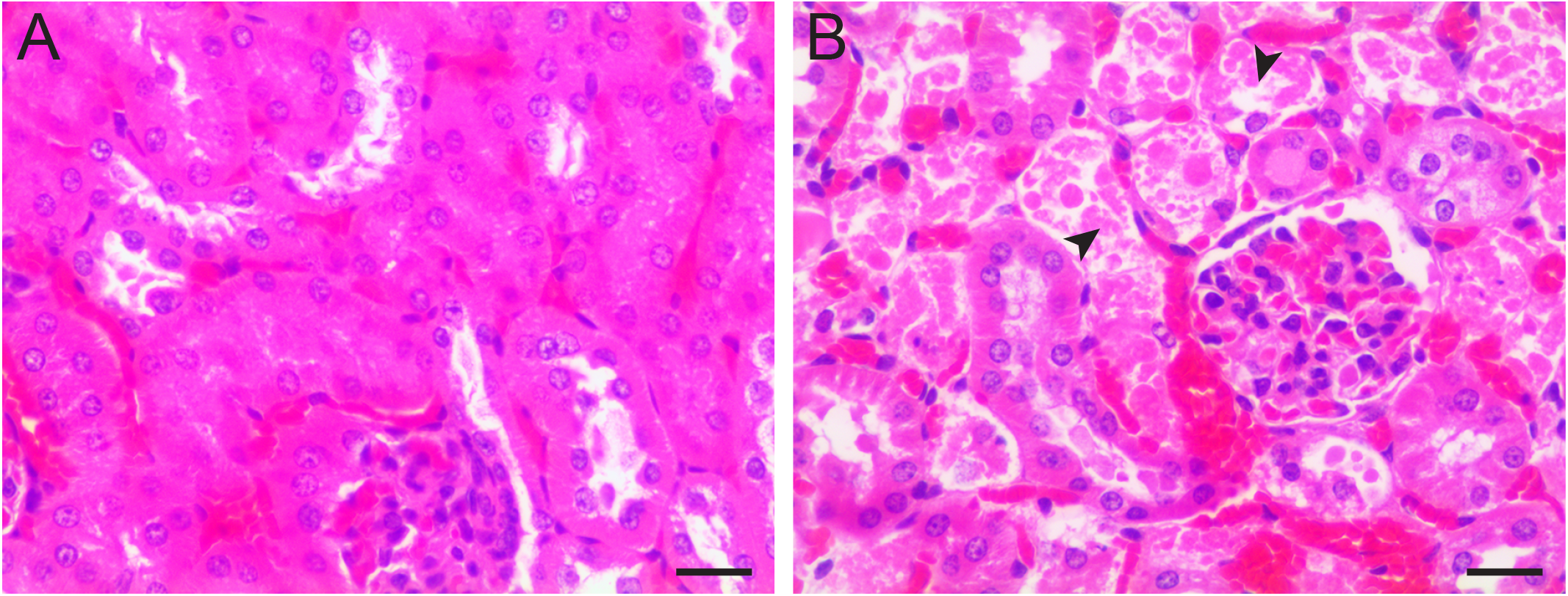
Delayed nephrotoxicity occurs at >10-fold higher doses of S-F than nourseothricin. **(A)** S-F dosing at 100 mg/kg without obvious histological abnormality in kidney. **(B)** Nourseothricin dosing at 10 mg/kg showing cellular necrosis and nuclear degeneration of proximal convoluted tubule epithelial cells (arrowheads). Glomeruli and distal tubules were spared. Tissue was harvested 3 days after dosing. Size bar = 20 μM.

### Murine thigh infection model

Therapeutic effect after a single dose of S-F was tested in the murine thigh model after infection with the pandrug-resistant Nevada CRE strain (AR-0636). For these studies, neutropenic mice were rendered mildly renal deficient with uranyl nitrate to more closely simulate human excretion kinetics (26). Tissue was harvested 24-hours post infection. In the absence of treatment, CFU increased > 2log_10_. In contrast, S-F at 50 mg/kg or 100 mg/kg led to a greater than 5log_10_ reduction in CFU with absence of detectable CFU in three of five mice treated with the higher dose (Fig. 6). Therefore, substantial in vivo bactericidal treatment effect was observed with a single dose of S-F at levels with minimal or without observable toxicity.

**Figure 6.**
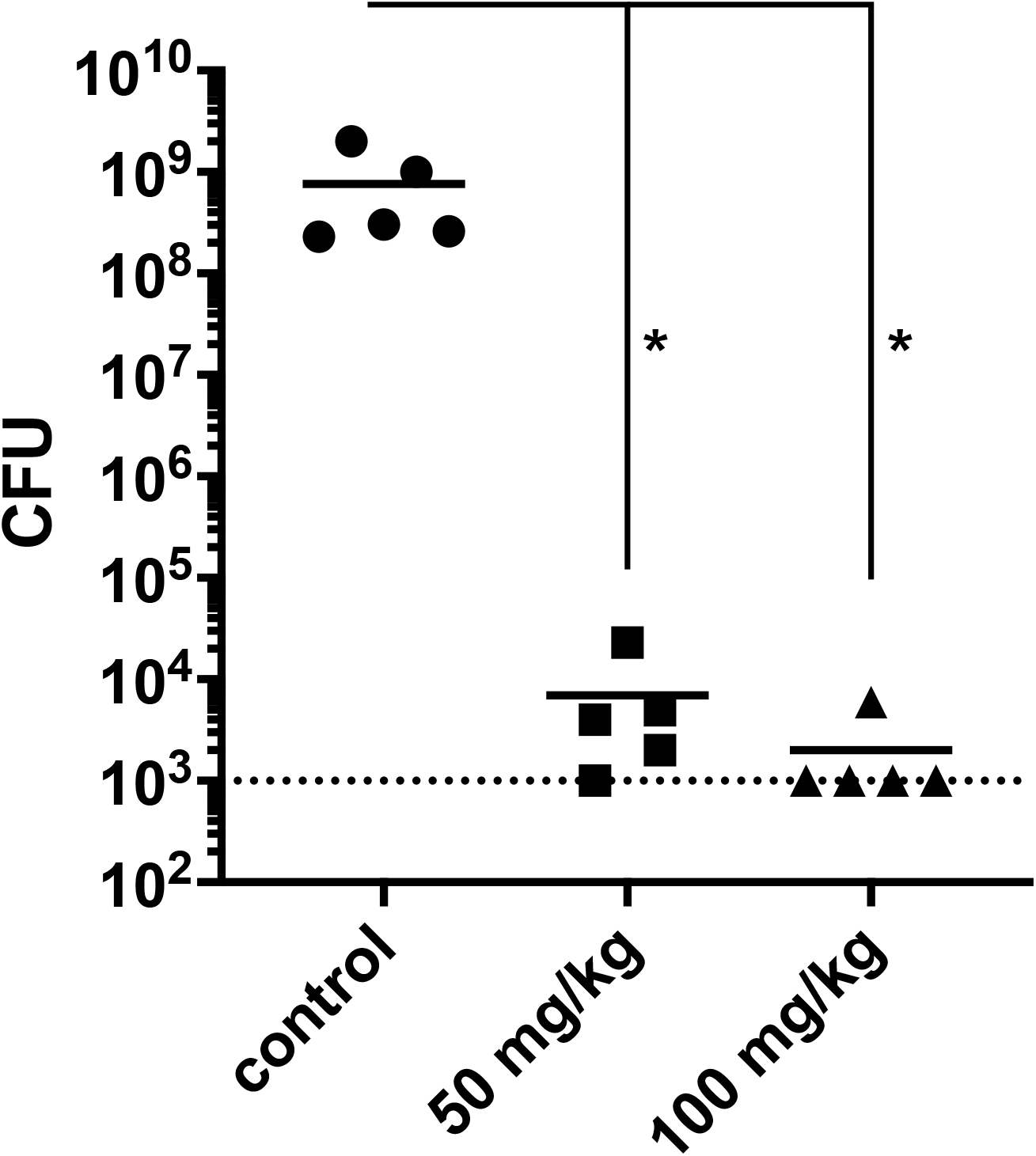
Murine thigh infection model. S-F demonstrated substantial therapeutic effect against pandrug-resistant *Klebsiella pneumoniae* Nevada AR-0636 at doses without observable or minimal toxicity. Dotted line is assay limit of detection. Data points for 5 mice per condition are shown. * designates significant difference from untreated controls using Kruskall-Wallis non-parametric test.

### Resistance studies suggest novel target

The antimicrobial activity of aminoglycosides is blocked by specific 16S rRNA methylases. Based on similarity in activity to aminoglycosides noted above, we considered the possibility that methylases that target the 16S rRNA A1408 position (e.g., NpmA) and block activity of all known 2-deoxystreptamine-based aminoglycosides including apramycin (5, 6), and methylases that target the 16S rRNA G1405 position (e.g., ArmA) and block activity of 4,6-disubstituted deoxystreptamine (DOS) aminoglycosides such as gentamicin, (but not apramycin), might interfere with activity of nourseothricin. Notably, in contrast to high level resistance to gentamicin and apramycin controls, activity of nourseothricin (and by inference its component streptothricins) was completely unaffected by cloned *npmA* and *armA* methylases expressed from a pBAD promoter under inducing conditions (see Table I). Furthermore, the single ribosomal operon *E. coli* strain SQ110 (27) with either A1408G or G1491A mutations in 16S rRNA helix 44 that conferred high level resistance to aminoglycosides such as gentamicin, tobramycin, kanamycin, neomycin and apramycin did not affect nourseothricin susceptibility (data not shown). These data suggested that the target and corresponding mechanism of action of streptothricins are distinct from traditional aminoglycosides.

**Table I.** Lack of effect of ribosomal 16S rRNA G1405 (ArmA) and A1408 (NpmA) methyltransferases on nourseothricin minimal inhibitory concentration (MIC in µg mL^-1^).

| Antibiotic | *armA*<sup>a</sup> | *npmA*<sup>a</sup> | vector control<sup>b</sup> |
| --- | --- | --- | --- |
| gentamicin | >256 | 16 | 0.125 |
| apramycin | 1 | > 256 | 1 |
| nourseothricin | 0.25 | 0.25 | 0.5 |

<sup>a</sup>genes cloned under inducible pBAD promoter (see Fig. S4). 1% arabinose added to MIC panels for induction.
<sup>b</sup>Vector control, same pBAD vector expressing an unrelated protein (LSSOrange) cloned in the same position as the *armA* and *npmA* methyltransferases, providing a measure of the susceptibility of the strains in the absence of any resistance gene.
MIC values represent mode for at least 8 separate determinations in two biological replicates.

To consider further the potentially unique interactions with the ribosome, we sought to identify high level nourseothricin resistance mutants in single ribosomal operon *E. coli*, thereby allowing isolation of characteristically recessive mutations in ribosomal RNA. Accordingly, we therefore plated concentrated bacterial suspensions of *E. coli* strain SQ110 on LB agar medium containing doubling dilutions of nourseothricin. We were able to isolate two classes of high-level nourseothricin resistance mutants obtained at a frequency of ∼3.8 × 10^−9^ only in the single *rrn* operon, SQ110, *E. coli* mutant strain, not in wt *E. coli* which has 7 *rrn* operons, implying mutations were recessive, as expected. Specifically, we obtained multiple independent mutants at 16S rRNA C1054A and C1054T with 32, 64 and 128 µg/mL nourseothricin selective concentrations, and A1196C and A1196G with 32 and 64 µg/mL selective concentrations. Both positions lie within helix 34 of the 16S rRNA. In total 54, 2, and 5 independently isolated strains containing C1054A, A1196C, and A1196G, respectively, were identified. In addition, C1054T was identified in 16 strains that proved non-viable on passaging from frozen stocks and were therefore not further analyzed. In Table 2, we show modal MIC values from three biological replicates from two randomly selected mutants of each type to rule out potential contributions of secondary mutations. C1054 and A1196 mutants conferred >256-fold and 16-32-fold increases in MIC for nourseothricin, respectively.

**Table 2.** Modal MIC values for Resistance Mutants Identified in Single rrn Operon *E. coli* Strain Identifies Unique Target in Helix 34 of Ribosomal 16S rRNA.

| strain | SQ110 | N1 | N8 | N9 | N11 | N12 | N13 |
| --- | --- | --- | --- | --- | --- | --- | --- |
| genotype | wt | C1054A | C1054A | A1196G | A1196G | A1196C | A1196C |
| nourseothricin | 1 | >256 | >256 | 32** | 16* | 64 | 64 |
| tetracycline | 2 | 4 | 8 | 4 | 2 | 4* | 4* |
| doxycycline | 2 | 8 | 8 | 4 | 4 | 2* | 2 |
| minocycline | 2 | 16 | 8 | 4 | 4 | 4* | 4 |

SQ110 contains only a single *rrn* operon parent strain. Random nourseothricin SQ110 resistance mutants were selected for by plating on high concentrations of nourseothricin. Mutations in 16S rRNA identified in these resistant strains are listed under genotype. MIC values are in $\mu\text{g/mL}$ determined after 40 hours of incubation, as single ribosomal RNA strains, especially mutants grew slowly, and differences between growth and inhibition could not be reliably distinguished at earlier time points. Modal MIC values from at least three biological replicates are shown. Unless otherwise state MIC range two doubling dilutions. \*The range of MICs obtained varied over three doubling dilutions.\*\*MIC range was 32-256 $\mu\text{g/mL}$ in four biological replicates.

Despite overlap of identified nourseothricin resistance mutations with the reported binding pocket of tetracyclines in helix 34 (28), biological data suggested that binding of nourseothricin and tetracyclines is distinct. Specifically, C1054A only conferred modest cross-resistance and A1196C/G conferring negligible if any cross-resistance to tetracycline, doxycycline, and minocycline (see Table 2). In addition, ribosomal protection proteins, for example, TetM, are believed to confer high level resistance to earlier generation tetracyclines through disruption of stacking interactions of these antibiotics with C1054 (29, 30). Although not performed with isogenic strains, we noted in a limited survey of *Staphylococcus aureus* isolates, that while tetM containing isolates were as expected highly tetracycline resistant, all isolates, whether containing TetM or not, remained highly susceptible to nourseothricin, suggesting that nourseothricin, unlike first and second generation tetracyclines, can bypass the mechanism of action of ribosomal protection proteins (see Table S3).

## DISCUSSION

We demonstrate that streptothricins, in particular S-F, S-D and the natural product mixture, nourseothricin, are highly active against contemporary carbapenem-resistant *E. coli, Klebsiella*, and *Enterobacter*. The activity spectrum is therefore highly relevant to a current and expected gap in gram-negative antimicrobial coverage.

Most of the existing description of streptothricin class antibiotic activity was from several decades ago and made use of unstandardized microbiological and physicochemical methods. In general, the natural product mixture was characterized as a whole without quantitative delineation of contributions from major constituent streptothricins. Instead, here we made use of modern preparative techniques and methods to characterize the activity of S-F and S-D in comparison with the natural product mixture currently available, gaining valuable insights into their respective properties.

We found that S-F maintained potent antimicrobial activity, although it was about six-fold less potent than S-D. Importantly, S-F showed at least 10-fold lower toxicity than S-D and the natural product mixture, nourseothricin, *in vitro* and *in vivo*. Notably, treatment with the nourseothricin natural product mixture was associated with proximal tubule kidney damage at relatively low doses (10 mg/kg), making nourseothricin or the likely even more toxic S-D component non-viable as therapeutics, consistent with prior observations. However, S-F appeared to have a much more favorable therapeutic ratio with the observation of substantial therapeutic effect to the point of single-dose sterilization of CRE infection within the experimental limit of detection without observable or minimal toxicity in single maximal tolerated dose histopathological analysis.

Biologically, streptothricins act similarly to aminoglycosides; specifically, nourseothricin demonstrated rapid, potent bactericidal activity (31). Correspondingly, streptothricins were also previously found to strongly induce translational miscoding (15), presumably poisoning the bacterial cell with junk protein.

It was therefore an original expectation that streptothricins would bind to the 30S ribosome in the same location as aminoglycosides. However, our resistance studies supported an alternative, unique mechanism of action. Specifically, resistance-conferring point mutations were found in helix 34 of 16S rRNA at C1054 and A1196. Notably, these point mutations did not impart resistance to any 2-deoxystreptamine (2-DOS) aminoglycosides tested (data not shown). Conversely, the 2-DOS aminoglycoside antibiotics are known to interact with helix 44 (residues 1400-1410 and 1490-1500) of 16S rRNA, forming a binding pocket near the ribosome decoding center (32-36). Correspondingly, the activity of 2-DOS aminoglycosides is blocked by methylation on residues 1405 and 1408, as well as by resistance-conferring point mutations at positions 1406, 1408, 1409, 1491, and 1495 (37-43). In contrast, neither methylation through action of ArmA and NpmA on 1405 and A1408, respectively, nor resistance mutations A1408G and G1491A affected nourseothricin activity. Of note, all of the forgoing mutations were created in single ribosomal operon strain, and could not be isolated in wild type (7 *rrn* operons) *E. coli* consistent with prior observations of the recessive nature of such aminoglycoside resistance mutations and highlighting the similar recessive nature of nourseothricin 16S rRNA resistance mutations.

The streptothricin mutations obtained are also distinct from those conferring resistance to other known antibiotics that interact with helix 34. This includes spectinomycin with resistance mutations isolated at positions A1191 and C1192 (37, 43-45), consistent with known contacts in structural studies (36), and negamycin, a pseudo dipeptide antibiotic, with U1060A, U1052G, A1197U resistance mutations (46, 47).

Tetracycline, has been reported to make hydrophobic contacts with 16S rRNA C1054 and 1196; however, our studies indicate only modest to negligible effects of C1054A and A1196C/G mutations on tetracycline, doxycycline, and minocycline MIC values (30). Our mutational data are also distinct from the previously described strong tetracycline resistance mutations, U1060A and G1058C (48-50). Therefore, the fundamental nature and importance of contacts appear different. Indeed, prior structural studies suggest interactions of tetracyclines primarily with the backbone of helix 34 (30); whereas the very strong mutational effects for nourseothricin (>256-fold increase in MIC for C1054A) suggest that the nucleoside moiety itself may be paramount for nourseothricin activity. Interestingly, TetM, which blocks activity of tetracycline through ribosomal protection and known contacts with C1054, appeared also not to confer resistance to nourseothricin.

Tetracyclines are thought to block binding of amino-acyl tRNA to the A-site codon through steric hindrance. As such, they do not cause miscoding, consistent with their largely bacteriostatic effects. In contrast, negamycin, though binding to similar area of helix 34, has been proposed to stabilize binding of near-cognate tRNAs to the A-site codon, thereby both inhibiting translation and promoting miscoding (47). Taken together, these observations suggest the mechanisms of action of streptothricins and tetracyclines are biologically distinct, while noting biological properties shared with negamycin, despite a distinct mutational resistance profile.

The specific nourseothricin resistance mutants in helix 34 obtained are noteworthy. Helix 34 has previously been implicated in ribosomal decoding, translocation, and termination functions (51). Intriguingly, C1054A was the first known ribosomal suppressor mutation of the UGA stop codon (52-54), supporting specific roles in termination and decoding. Intriguingly, both C1054, lying within a conserved bulge in helix 34, and A1196 are conspicuously unpaired in secondary RNA structure and were observed to have stacking interactions with one another (55), potentially providing a combined nidus for streptothricin interaction. C1054 has been noted to lie in proximity to the A-decoding site, and its close neighbor, U1052, in crosslinking studies was found to interact with the third nucleotide of the A-site codon in mRNA (56, 57). Elongation factor G (EF-G) was also previously found to specifically protect C1054 and A1196 from chemical modification (51) and structural studies show interaction of EF-G, domain IV with C1054 (29, 51). Although the forgoing associations place streptothricin in proximity to key mediators of translational fidelity and processivity, further understanding of streptothricin mechanism of action will likely await dynamic structural biology studies.

In summary, we present data for compelling bactericidal activity of S-F against contemporary multidrug-resistant CRE pathogens with *in vivo* confirmation of efficacy against an emblem of Gram-negative antibiotic resistance. We also find evidence for unique mechanism of action targeting 16S rRNA helix-34 distinct from 2-DOS aminoglycosides and tetracyclines. We therefore believe that further early-stage exploration of the historic scaffold is warranted with the goal of identification of analogs with potential for therapeutic development.

## MATERIALS AND METHODS

### Microbial strains

CRE strains were from the FDA-CDC Antimicrobial Resistance Isolate Bank (AR-Bank) including the Nevada Strain (AR-0636) and the BIDMC clinical collection sequenced as part of the CRE genomics project (58). Strains are listed in Table S1. The single ribosomal operon strain, *E. coli* SQ110, was from the Coli Genetics Stock Center (Yale University, New Haven, CT) and was originally created and described by Squire and colleagues (27).

### Streptothricin purification

Purification of separate streptothricin compounds from nourseothricin sulfate (Gold Biotechnology, St. Louis, MO) was performed through modification of a previously reported method (17). A glass column (150 cm x 2.4 cm) was packed with Sephadex LH-20 size exclusion gel (GE Healthcare, Chicago, IL) using a mobile phase of 10% methanol / H_2_O. The flow rate was adjusted using compressed air to 0.6 mL / min. Purifications were run in batches of approximately 300 mg of nourseothricin sulfate which was diluted in 0.6 mL of H_2_O and loaded dropwise directly onto the top of the column. A mobile phase of 10% methanol / H_2_O was used for elution and fraction sizes of 3 mL were collected. Fractions testing positive for the ninhydrin stain were analyzed for purity by LC-MS. Pure streptothricin D began eluting after approximately 120 mL of mobile phase, followed by mixed fractions of streptothricin D / E / F, and finally pure streptothricin F. Pure fractions for streptothricin D were combined, frozen, and lyophilized to give a powdery, off-white solid. Pure fractions for streptothricin F were combined, frozen, and lyophilized to give a powdery, off-white solid. Additional experimental details including ^1^H- and ^13^C-NMR spectra, mass spectrometry, as well as elemental analysis data of streptothricin F and streptothricin D can be found in the supporting information.

### MIC testing

The Clinical Laboratory and Standards Institute (CLSI) broth microdilution reference method was used for MIC testing (59). Bacterial inocula were prepared by passaging previously frozen bacterial strains on sheep blood agar plate, culturing for 18-24 hours, except where noted, and suspending isolated colonies to 0.5 McFarland (∼1×10^8^ CFU/mL) in 0.9% NaCl solution. This suspension was diluted 1:300 into cation-adjusted Mueller-Hinton broth (BD Diagnostics, Franklin Lakes, NJ) to achieve a final inoculum concentration of ∼5×10^5^ CFU/mL in 100 µL well volumes in round-bottom, 96-well plates (Evergreen Scientific, Los Angeles, CA). Doubling dilutions or antimicrobials were dispensed using the D300 digital dispensing method for final doubling dilution concentrations ranging from 0.125 to 256 µg/mL (60-62).The inkjet printing methodology was previously extensively validated and found to be as accurate and more precise the manual dilution techniques described in CLSI methodology (62). MIC values were determined after incubation for 16-20 hours.

### Time-kill studies

Time-kill studies were performed as previously described (63). At time 0, antibiotics and inoculum were combined in cation-adjusted Mueller-Hinton broth (BD Diagnostics, Franklin Lakes, NJ) at various antibiotic concentrations based on multiples of each isolate’s MIC as determined through broth microdilution assays. Each tube was incubated at 35°C in ambient air on a platform shaker. Sterility, treatment, and no antibiotic growth controls were plated at 0-, 1-, 2-, 4-, 6-, and 24-hour time points, respectively. At each time point, the number of viable bacteria were quantified using the drop plate method (64) as a substitution for the CLSI spread plate method. Specifically, 10 µL drops of serial ten-fold serial dilutions of bacterial suspension were plated. As previously described (64), only drops with 3 to 30 colonies were counted. The limit of detection for this assay was conservatively set at 300 CFU mL^-1^ taking into account Poisson distribution considerations.

### *In vitro* translation assays

Prokaryotic and eukaryotic nanoluciferase reporter constructs were constructed as follows: To construct the *in vitro* bacterial nanoluciferase (Nluc) transcription-translation expression system, a DNA fragment containing the 516-bp (see Table S3) from pNL1.1 Promega, (Madison, WI), including the NLuc open reading frame and Shine-Delgarno sequence, with addition of a T7 promoter sequence (TAATACGACTCACTATAGG), was synthesized as a gBlock by IDT (Integrated DNA Technologies, Coralville, IW). The T7_Nluc fragment was then amplified using primer pairs, T7Nluc-F and T7NLuc-R (see Table S5), and ligated into the pCR XL TOPO vector (Invitrogen, Carlsbad, CA) to create pCR XL TOPO-T7NLuc (Fig. S4A), and transformed into E. coli NEB-5α for plasmid preparation.

To construct the eukaryotic Nluc *in vitro* transcription-translation system, then Nluc open reading frame without the methionine initiation codon was amplified with primers, Nluc_BamHI-F and Nluc_PstI-R (see Table S5). Amplicon was digested with BamHI and PstI, cloned into similarly digested pT7CFE1-NFTag in frame with leading sequence and downstream of vector IRES and T7 promoter sequences (#88865, Thermo Fisher, Waltham, MA) to create pT7CFE1-NLuc (Fig. S4B), and transformed into *E. coli* NEB 5-alpha for plasmid preparation.

Nourseothricin, S-D, and S-F were distributed to a 384-well black plate (PHENIX Research, Swedesboro, NJ) at concentrations of 0-52 μM for bacterial *in vitro* expression or 0-820 μM for eukaryotic *in vitro* expression. For bacterial *in vitro* expression, we added *E. coli* S30 Extract System for Circular DNA (Promega, #L1020) to each well with introduction of 100 ng of pCR XL TOPO-T7NLuc per reaction making use of the upstream vector, pLac promoter rather than the cloned T7 promoter for expression in the current studies. After incubation at 37°C for 60 minutes, PBS buffer having 1,000 volume diluted Nluc-specific furimazine substrate (Nano-Glo Luciferase Assay System, Promega) was added to each well, and luminescent signal was detected with an Infinite M1000 Pro (TECAN, Morrisville, NC) plate reader. Eukaryotic *in vitro* Nluc expression was performed with TnT-T7 Quick Coupled Transcription/Translation System (# L1170, Promega) with 100 ng of pT7CFE1-NLuc plasmid DNA, TnT master mix, and methionine distributed to test wells per manufacturer’s instructions. After incubation at 30°C for 80 minutes, Nluc signal was detected as described above.

### Eukaryotic cell cytotoxicity

J774A.1 mouse macrophage and LLC-PK1 porcine renal tubule epithelial cell lines were plated in 384-well tissue culture dishes with white walls and clear bottoms (Greiner, Monroe, NC, catalog #781098) at ∼7.0 × 10^5^ cells/cm^2^ and ∼7.0 × 10^4^ cells/cm^2^, respectively, in M199 medium lacking phenol red supplemented with 3% porcine serum and a final concentration of 125 nM SYTOX Green. Approximately one day after plating (at 50%-75% confluence), the cell lines were treated with two-fold doubling dilutions of nourseothricin (Gold Biotechnology), S-D, and S-F, dispensed with the HP D300 digital dispensing system (HP, Inc., Palo Alto, CA) using Tween-20 as the surfactant. Final surfactant concentration was adjusted to 0.0004% in all wells. Microplates were vortexed for 30 seconds to ensure thorough mixing within the wells prior to incubation. SYTOX Green fluorescence was read with a TECAN M1000 multi-mode reader using excitation 485 ±7 nm and emission 535 ±10 nm settings at indicated time points with otherwise continuous incubation at 37°C with 5% CO_2_ for 5 days.

### Maximum tolerated dose (MTD) determination

CD-1 (ICR) female mice weighing 25-30 g each were purchased from Charles River Laboratories, Inc. (Kingston, NY). Mice were injected intraperitoneally (IP) with 0, 10, 20, 50, 100, 200, 400, mg kg^-1^ of purified streptothricin-F or nourseothricin. On day 3 post dosing, mice were euthanized and kidneys fixed in 10% neutral phosphate-buffered formalin overnight, embedded in paraffin, and 10 μm tissue sections stained with hematoxylin and eosin at an institutional core facility using standard procedures as described previously (65).

### Neutropenic mouse thigh model

Mice were pretreated with cyclophosphamide and uranyl acetate and infected by injection of 10^6^ CFU of the CRE Nevada strain AR-0636 suspended in 100 μL PBS into the thigh muscle (66). Mice were then injected with streptothricin-F administered subcutaneously at the scruff of the neck (i.e., distant from the thigh) two hours post infection and euthanized after 24 hours. Thigh tissue was excised and homogenized using a disposable tissue grinder (Fisher Scientific) in 1 mL cation-adjusted Mueller-Hinton broth. Homogenates were serially diluted, plated on Mueller-Hinton agar plates (Remel, Lenexa, KS), and incubated at 37°C overnight for CFU enumeration.

### Cloning of 16S rRNA methylases

The genes for the 16S rRNA G1405 and A1408 methylases, *armA* and *npmA*, were codon optimized for *E. coli* based using the ITD Codon Optimization Tool (https://www.idtdna.com/pages/tools/codon-optimization-tool,), synthesized as a gBlock (IDT, Coralville, IA), and used to replace the LSSmOrange open reading frame in the arabinose-inducible vector, pBAD-LSSmOrange (67), a gift from Vladislav Verkhusha (Addgene plasmid # 37129; http://n2t.net/addgene:37129; RRID:Addgene_37129) by HiFi assembly (New England Biolabs, Beverely, MA) using primers, pBAD_F, pBAD_R, and armA_F and arm A_R; and npmA F and npmA_R, respectively (Table S5) Assembled vectors (Figs. S4C, S4D) were transformed into NEB 5-alpha (New England Biolabs), selecting for ampicillin and gentamicin resistance. MIC assays were performed in the presence of 1% arabinose for resistance gene induction.

### Mutational Resistance Studies

Single ribosomal operon strain, SQ110, was grown overnight in CAMHB at 37°C and resuspended in one-fifth volume in fresh medium. Thereafter, 200 uL of the suspension was plated on selective LB agar plates containing nourseothricin (GoldBio, St Louis, MO) at multiple two-fold doubling dilution concentrations and incubated for 48-72 hours at 35°C in ambient air until mutant colonies became visible. Selected colonies were passaged on plates with matching concentrations of nourseothricin for further analysis. Aminoglycoside resistance mutants were isolated in a similar manner selecting on apramycin.

Genomic DNA from nourseothricin resistant strains was extracted using the Wizard Genomic DNA Purification Kit (Promega, Madison, WI) from 3 mL of overnight culture in CAMHB containing NAT [32 µg/mL, 64 µg/mL, and 128 µg/mL]. Subsequent genomic DNA was amplified using primers F16S+23S, R16S+23S, and 16SR (Table S5), with primer 16SR as previously described (27)). All amplification reactions were performed using Q5 high-fidelity DNA polymerase, Q5 reaction buffer, and DNTPs from New England Biolabs (Ipswich, MA) using a melting temperature of 68°C, an annealing temperature of 65°C, and an extension time of two minutes and 45 seconds. PCR products were purified using the QIAquick Purification Kit. Sanger sequencing was performed to identify mutations.

## ACKNOWLEDGEMENTS

This work was supported by R21 AI140212 to R.M. and J.E.K, and R01AI157208 to J.E.K., R.M., and E.Y. K.P.S. and K.E.Z. were supported by the National Institute of Allergy and Infectious Diseases of the National Institutes of Health under award numbers F32 AI124590 and T32 AI007061, respectively. The content is solely the responsibility of the authors and does not necessarily represent the official views of the National Institutes of Health. The HP D300 digital dispenser and TECAN M1000 were provided for our use by TECAN (Morrisville, NC). Tecan had no role in study design, data collection/interpretation, manuscript preparation, or decision to publish.

